# Analysis of the structural variability of topologically associated domains as revealed by Hi-C

**DOI:** 10.1101/498972

**Authors:** Natalie Sauerwald, Akshat Singhal, Carl Kingsford

## Abstract

Three-dimensional chromosome structure plays an integral role in gene expression and regulation, replication timing, and other cellular processes. Topologically associated domains (TADs), one of the building blocks of chromosome structure, are genomic regions with higher contact frequencies within the region than outside the region. A central question is the degree to which TADs are conserved or vary between conditions. We analyze a set of 137 Hi-C samples from 9 different studies under 3 measures in order to quantify the effects of various sources of biological and experimental variation. We observe significant variation in TAD sets between both non-replicate and replicate samples, and provide initial evidence that this variability does not come from genetic sequence differences. The effects of experimental protocol differences are also measured, demonstrating that samples can have protocol-specific structural changes, but that TADs are generally robust to lab-specific differences. This study represents a systematic quantification of the key factors influencing comparisons of chromosome structure, suggesting significant variability and the potential for cell-type-specific structural features, which has previously not been systematically explored. The lack of observed influence of heredity and genetic differences on chromosome structure suggests that factors other than the genetic sequence are the drivers of this structure, which plays such an important role in human disease and cellular functioning.

## Background

While it is recognized that the three-dimensional structure of the chromosome is an integral part of many key genomic functions, we lack a full understanding of the variability of this structure across biological sources or experimental conditions. Changes in chromosome structure at specific genomic regions and under certain conditions have been implicated in a variety of human diseases and disabilities, including many cancers [26, 28, 18, 19], deformation or malformation of limbs during development [25], and severe brain anomalies [41]. In healthy cells, genome shape is heavily linked to key processes such as gene regulation and expression [8, 5, 22, 14, 36], replication timing [37, 29, 33, 2], and DNA accessibility and nuclear organization [44, 34, 6]. Despite the clear importance of these structures, there has been no systematic study of the expected variation of topologically associated domains (TADs) genome-wide.

Features of genome-wide, three-dimensional chromosome structure can be measured by Hi-C [24], a high-throughput variant of the chromosome conformation capture protocol [9]. The experiment involves cross-linking and ligating spatially close genomic segments, then aligning them back to the genome to find their genomic positions. The output of this experiment is a matrix in which the rows and columns represent segments of the genome along a chromosome, and each matrix entry records the pairwise interaction frequency of the genome fragments of the associated row and column. These values reflect 3D proximity, quantifying the frequency of contact between every pair of genomic segments.

A hierarchical architecture has emerged from these Hi-C matrices, in which chromosome structure is composed of several different scales of components, from multi-megabase compartments to sub-megabase TADs and sub-TADs [15, 4]. TADs represent chromosomal regions with significantly higher interaction frequency among segments within the TAD than with those outside of it [11]. TADs are considered to be a primary structural building block of chromosome architecture [13], and several methods have been developed to computationally identify them [11, 35, 7, 43, 15, 31].

One challenge in the interpretation of TADs is that we have little understanding of the variability of TAD structures under different conditions. While some work has compared aspects of Hi-C data quality, TADs in particular were not considered [46]. No other Hi-C study has used more than 23 samples from different conditions, and even large data repositories such as ENCODE and the 4D Nucleome contain no more than 30 human Hi-C samples on their own. As more Hi-C data has become available recently, it is now possible to perform a substantial analysis of the relative consistency or variability of TADs across a variety of human cell conditions, by combining Hi-C samples from many studies and resources. Previous work has suggested that TADs are largely conserved across human cell types and possibly even species, however the degree of this conservation is unclear and has been tested in only small sets of samples [11, 35].

An initial method to compare TADs between cell types was previously applied to compare normal versus cancer human cell types [38], but that study did not investigate other potential sources of TAD variability and only compared 23 different cell or tissue types. We instead systematically quantify several sources of variability that have not been previously studied, using over three times as many different cell conditions, and three metrics.

We quantify the influence of both technical and biological variation on TAD structures across several experimental and biological conditions in the first study of over 100 Hi-C experiments. We observe that 10–70% of combined TAD boundaries differ between replicates, regardless of sequencing depth or contact coverage, pointing to a potentially dynamic or disordered arrangement. Across 69 samples of different cell lines and tissue types, we observe ~20–80% unshared TAD boundaries, suggesting that there can be fairly large differences in TAD sets across biological conditions, in contrast to previous claims of extensive TAD conservation [11, 35, 39]. We find that samples of the same cell or tissue type have elevated structural similarities, suggesting that biological function is a key driver of structural similarity. Though it is commonly believed that TADs do not vary much across cell types and possibly even species, we observe significant TAD variation across human cell and tissue types. By analyzing the structural similarity of sets of parents and their children, as well as tissue samples taken from different individuals, we observe that the genetic sequence differences between individuals and the genetic sequence similarities between parents and their children have little impact on TAD structural similarity. Of the possible sources of technical variation considered in this work, the choice of *in situ* (in nucleus) ligation versus dilution (in solution) ligation protocols has the greatest influence on Hi-C and TAD structures. In contrast, we demonstrate that Hi-C measurements and corresponding TAD calls are robust to other technical differences such as the choice of restriction enzyme and lab-specific variations.

## Results

### Data and comparison measures

We analyzed 69 publicly available human Hi-C samples (137 including replicates) from nine different studies [11, 35, 42, 10, 21, 12, 47, 17, 39] covering 52 unique biological sources and a variety of experimental conditions (Table 1). All samples were processed from raw reads (.fastq files) to normalized Hi-C matrices at 100kb resolution through the HiC-Pro pipeline [40]. TADs were computed on all resulting Hi-C matrices using Armatus [15].

**Table 1:**
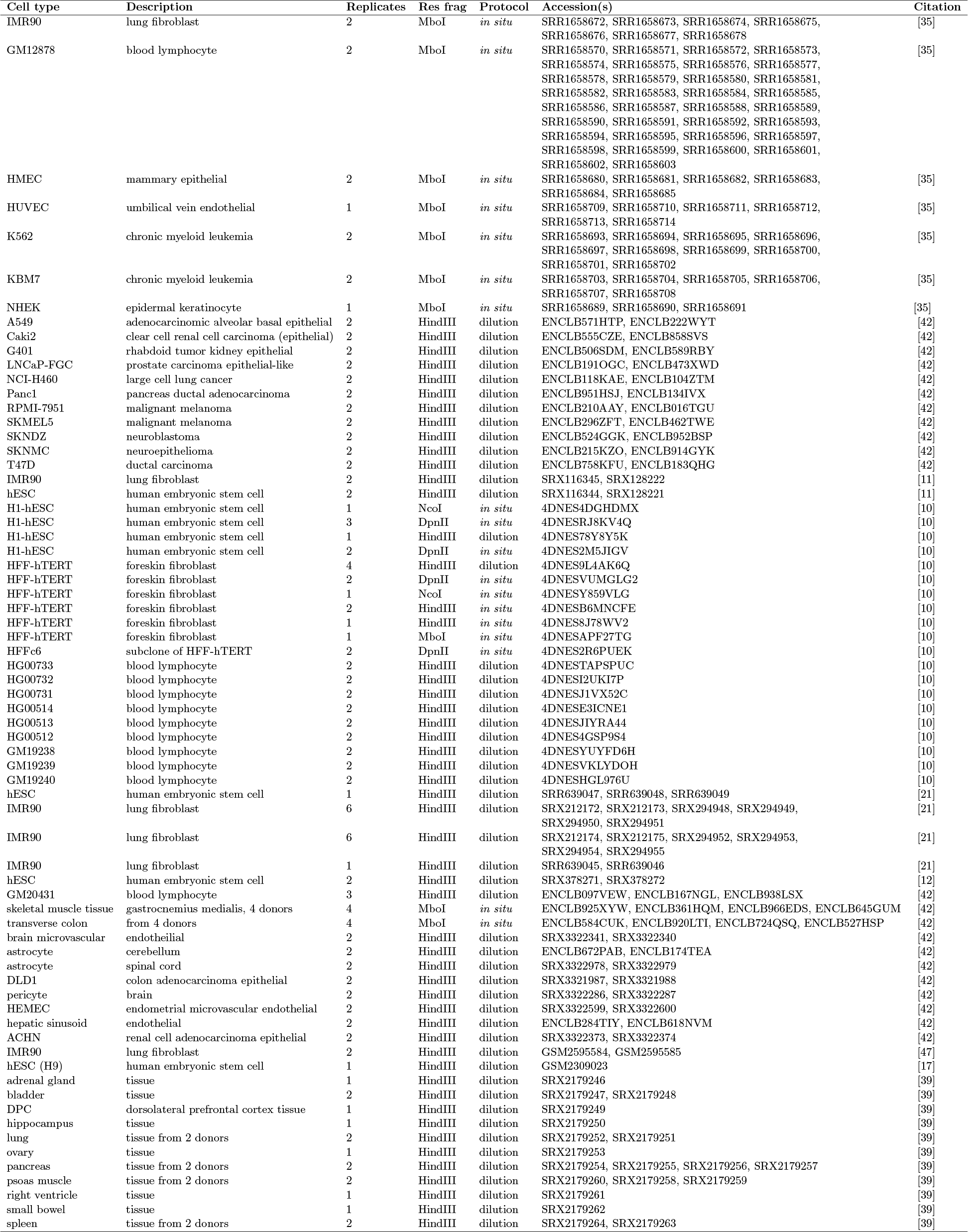
All Hi-C data used in this study

Pairwise similarity of these 69 samples (2346 pairs plus 83 replicate pairs) was quantified with three different similarity measures. HiCRep [45] computes a statistical similarity score on Hi-C matrices directly. The Jaccard Index (JI) quantifies the similarity between two sets, in this case defined as sets of TAD boundaries, returning a value that represents the fraction of TAD boundaries shared between the two samples. TADsim is a measure adapted from a method that identifies structurally similar regions between two TAD sets [38]; it quantifies the fraction of the genome covered by similar TAD structures using the variation of information (VI) metric [27]. See the Methods section for details on each comparison measure.

Distributions of similarity values are compared using the Mann-Whitney *U* test, which is a non-parametric statistical test quantifying the probability that two independent samples were drawn from populations with the same underlying distribution. More specifically, the alternative hypothesis for this test is that one distribution is greater than the other. A low *p*-value suggests that one of the two distributions is likely drawn from a distribution with a higher median than the other; we will use this to test whether various conditions reflect elevated structural similarity, relative to a background or other comparison distribution.

### Structural similarity of replicate samples

By quantifying the similarity of all 83 replicate pairs in our dataset, we find that the TAD sets of replicate pairs are significantly more similar than those of non-replicate pairs (Figure 1a–c, *p* < 10^−20^ for all comparison measures), in contrast to previous work that suggested much lower TAD reproducibility between replicates [16]. We note that this discrepancy can be explained by the fact that Forcato *et al.* [16] used a different pre-processing method which results in many fewer aligned reads than HiC-Pro and therefore significantly fewer Hi-C contacts, decreasing the reproducibility they observe. Using the same data, processed instead with HiC-Pro, at the same resolution and the same Armatus parameters gives JI values consistent with those reported here.

**Figure 1:**
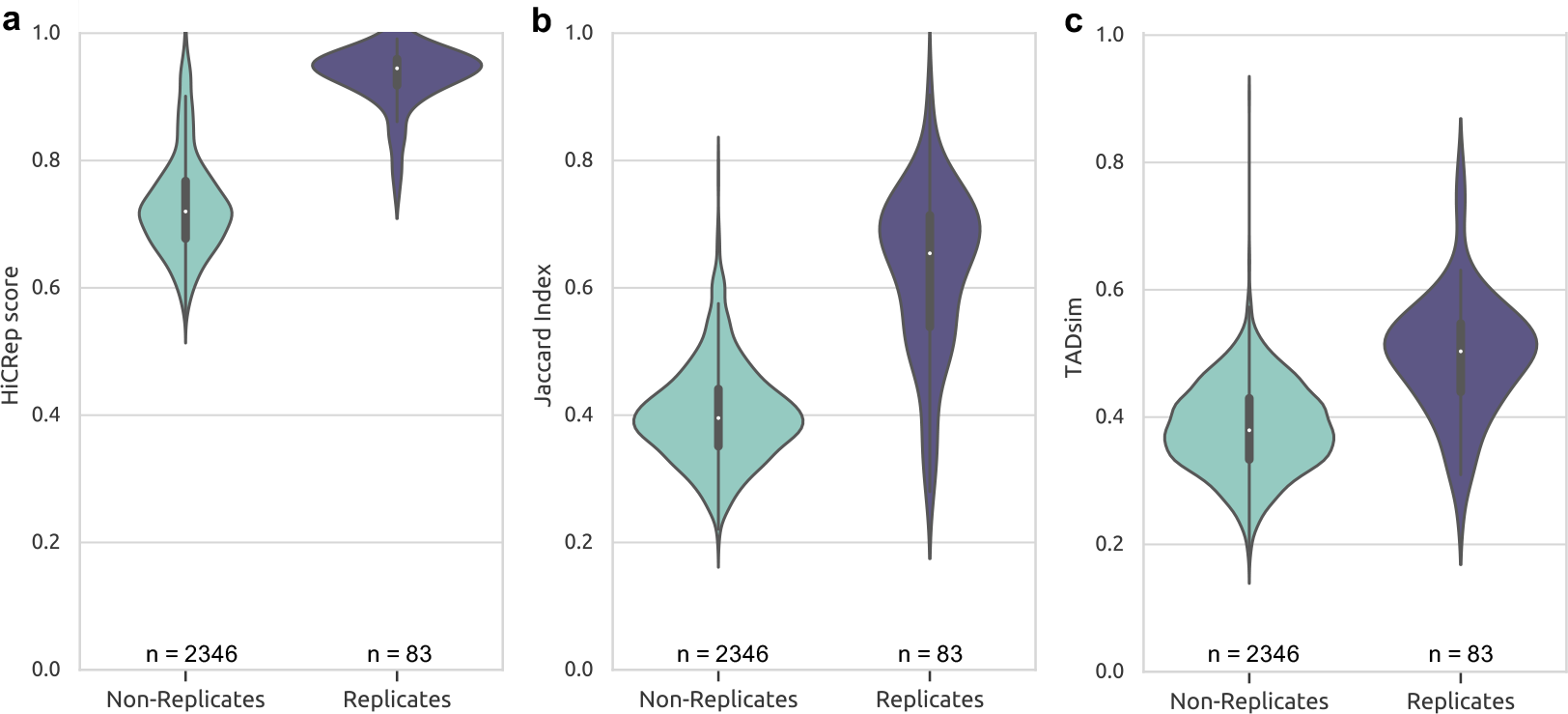
Hi-C and TAD reproducibility. The violin plots show distributions of **a:** HiCRep, **b:** Jaccard Index, and **c:** TADsim values on pairs of either replicates or non-replicates. All of these plots show a statistically significant (*p* < 10^−20^) elevation of similarity among replicate pairs, demonstrating that both Hi-C matrices and TADs are reproducible.

Among the samples studied in this work, replicates share an average of 62.77% of their combined TAD boundaries, which is consistent with other previous studies on different data using different methods (Dixon *et al.* [11]: 62.28%, 73.73% and Rao *et al.* [35]: 61.88%). This leaves almost 40% of TAD boundaries that vary across replicates. Between non-replicate pairs, almost 60% TAD boundaries are not shared on average, which contradicts the common notion that TADs are highly conserved between human cell types. This variability could not be explained by limitations of sequencing depth, as we found that reproducibility is only weakly correlated with sequencing coverage (see Figure S5). This points to a dynamic or disordered structure, as suggested by a recent imaging study [32], and a much higher level of TAD variation than previously thought.

### Variability across tissues and individuals

The chromosome structure of tissue samples has not been as widely studied as that of cell lines, but these structures may provide valuable insight into tissue-specific genome spatial organization. Among our set of 69 Hi-C experiments, 13 different human tissues are represented, and there are 16 pairs of the same tissue type taken from different donor individuals. The similarity values of the chromosome structures of these pairs are statistically indistinguishable from those of replicate samples (Figure 2a–c; HiCRep: *p* = 0.4792, JI: *p* = 0.1300, TADsim: *p* = 0.09559). There is much less variation across individuals than across tissue types (Figure 2a–c; HiCRep: *p* = 1.5577 × 10^−6^, JI: *p* = 3.876 × 10^−6^, TADsim: *p* = 1.017 × 10^−7^), suggesting that individual genetic differences have less influence on chromosome structure than the biological function of the sample.

**Figure 2:**
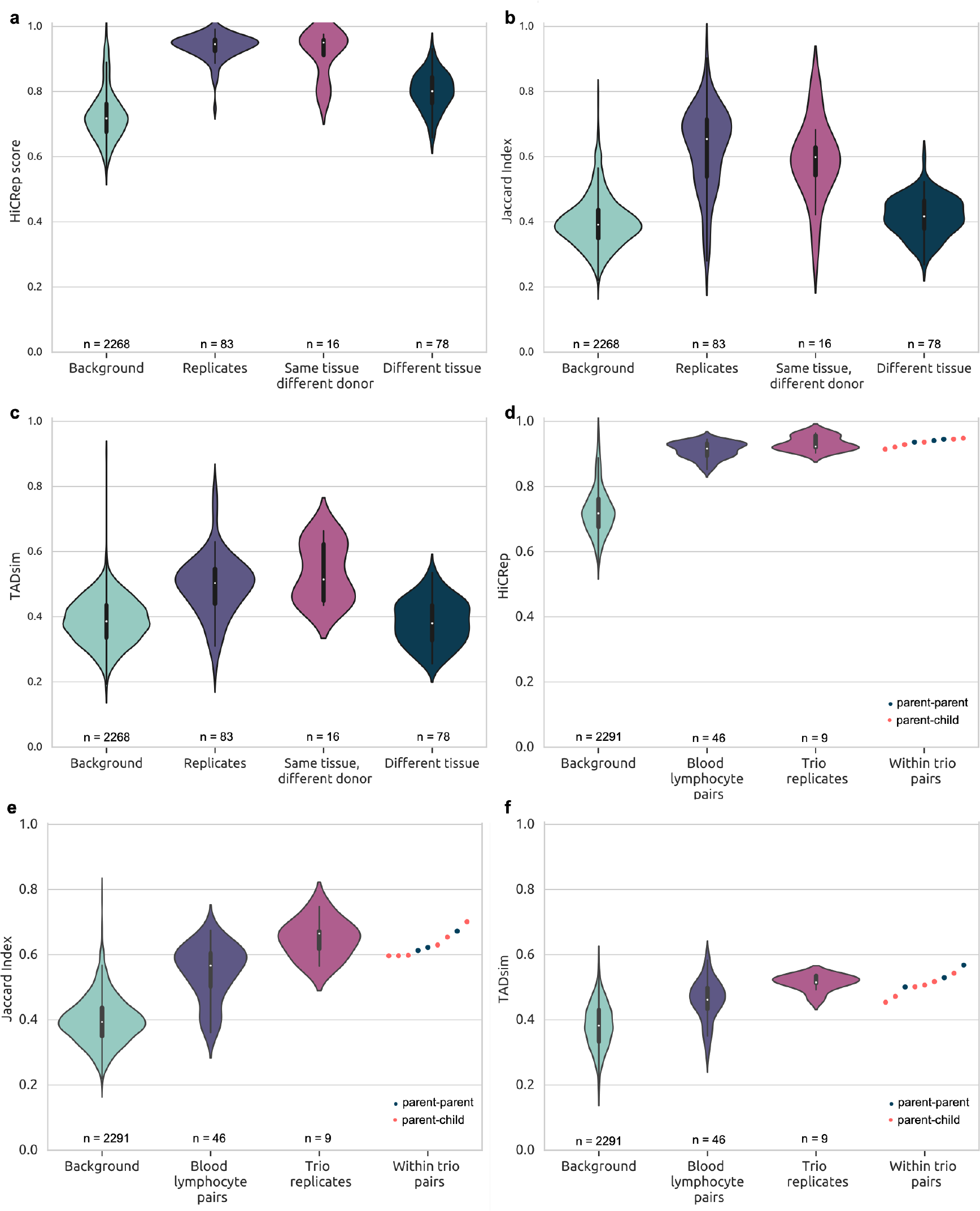
Biological sources of TAD variation. **a,b,c:** Comparisons between and within tissue samples using **a** HiCRep, **b** JI and **c** TADsim. Each figure shows four violin plots representing distributions of similarity values of the background (non-replicate pairs), replicates, pairs from the same tissue type but different donor individuals, and pairs from different tissue types. **d,e,f:** Comparisons with three trio samples of blood lymphocyte cells using **d** HiCRep, **e** JI and **f** TADsim. The background distribution consists of all non-replicate pairs, and the blood lymphocyte pair distribution shows all similarity values of two blood lymphocyte samples outside of the trios. The trio replicates refer to the similarity values of the replicate pairs from within each individual of the trio samples. The scattered points on the right side of each figure represent all within-trio comparisons, colored by family relationship.

Our analysis suggests that around 40% of TAD boundaries are shared between different tissue types, consistent with the findings of Schmitt *et al.* [39]. While this is significantly more than expected given random TAD boundary locations, it leaves room for large differences in the TAD sets of different tissue samples. In order to determine whether TAD structure is more similar across tissues than across cell lines, we compared the similarities between tissue types to the background distribution consisting of all non-replicate pairs with at least one cell line. Two of our three measures suggest that there is elevated conservation between tissues compared with cell lines, but the TADsim results suggest these values are statistically similar (Figure 2a–c, HiCRep: *p* = 2.120 × 10^−19^, JI: *p* = 0.004806, TADsim: *p* = 0.4235). The average JI value between tissue samples of 41.6% implies that while there is a significant level of similarity among chromosome structures of different tissue types, close to 60% of TAD boundaries vary between different tissue samples. This level of variability between tissue types may indicate the existence of tissue-specific structural features, rather than significant conservation of TADs between tissue types.

### Family relationships do not seem to influence TAD similarity

Hi-C measurements from blood lymphocyte cells of three parent-parent-child triplets (trios) permit a glimpse into the heritability of chromosome structure. We find that unrelated individuals (parents) share just as much structural similarity as each parent and their child (Figures 2d–f). We therefore see no evidence that chromosome structure is determined by genetic similarity, at least in blood lymphocyte cells. The similarity values within trios are generally much higher than the background of non-replicate comparisons, however they are similar to the distribution of pairs of blood lymphocytes, so this is likely a result of the shared cell type rather than genetic similarity. As with the tissue data, the biological source (cell or tissue type) seems to be a much stronger driver of structural similarity than genetic similarity.

### Variations across Hi-C protocols

In order to investigate technical sources of variation, we compare several common variations in the Hi-C protocol, and test whether they affect the similarity of the TADs that are identified. There are two main protocol variants that differ in the cross-linkage step. *In situ* Hi-C [35] (also termed “in nucleus Hi-C” [30]) involves cross-linking the DNA within the nucleus, while dilution Hi-C (or “in solution Hi-C”) performs cross-linking in a dilute solution. Each protocol also requires the choice of a restriction enzyme, which could be any of four common options: HindIII, MboI, NcoI, and DpnII. While there has been some study of the differences in Hi-C data resulting from *in situ* and dilution protocols [35, 30], the influence on TAD sets and the effect of the restriction enzyme had not been systematically studied previously.

### *In situ* and dilution Hi-C reproducibility

Across all replicate pairs (12 in situ, 71 dilution), the intra-chromosomal Hi-C matrices of *in situ* replicates are statistically significantly more similar than dilution replicates (Figure 3a, *p* = 1.180×10^−5^). However, the TAD sets of *in situ* replicates only show statistically significantly higher similarity than those of dilution replicates under the JI measure (Figures 3b,c; JI: *p* = 0.02703, TADsim: *p* = 0.1547). TADs capture only relatively short-range interactions, and it therefore appears that the difference between *in situ* and dilution Hi-C is not as significant a factor in TAD reproducibility as in overall Hi-C matrix reproducibility. It has been previously shown that *in situ* Hi-C matrices are more reproducible than dilution Hi-C matrices [35, 30], specifically with respect to long-range and inter-chromosomal contacts.

**Figure 3:**
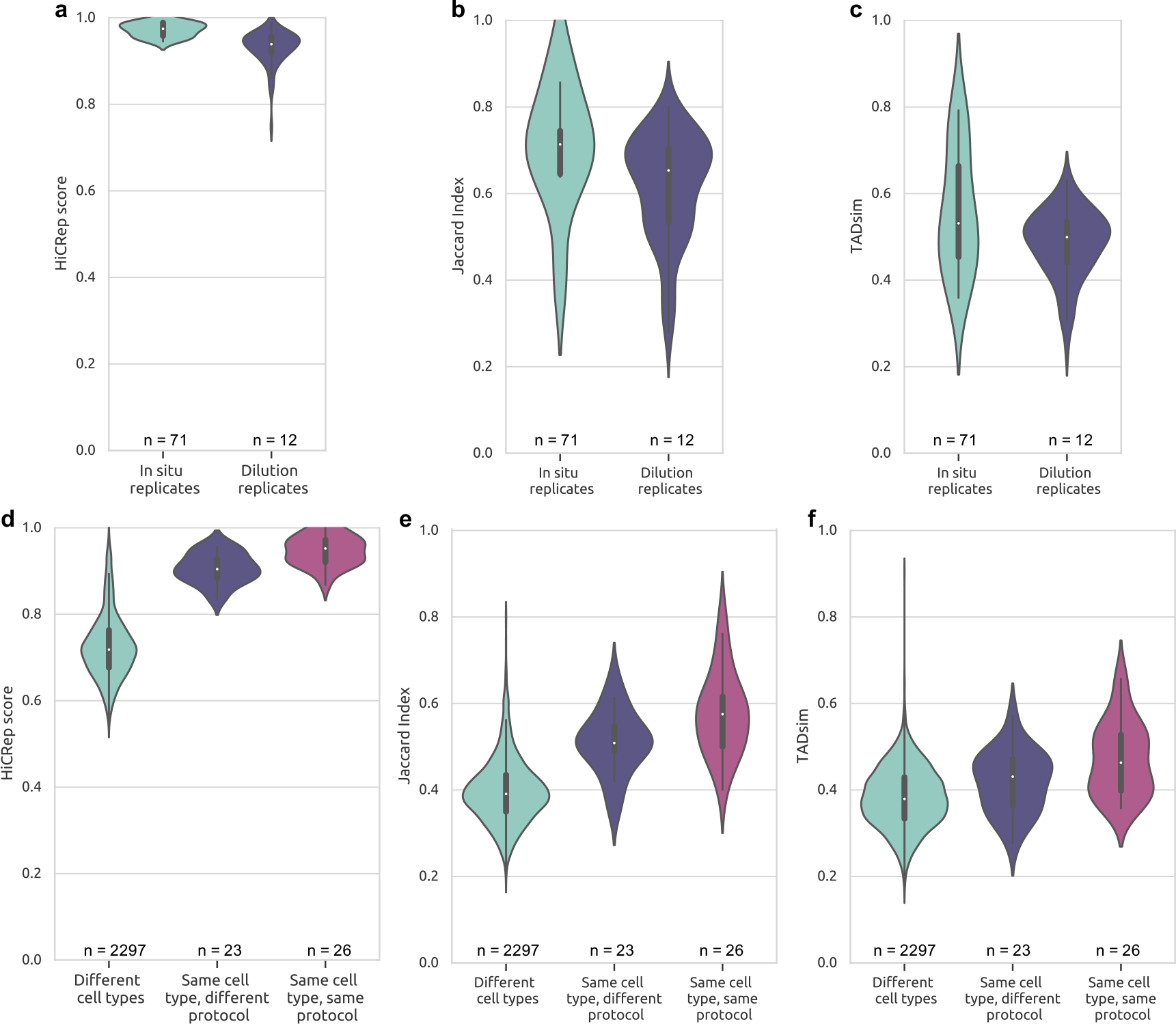
Comparing Hi-C samples generated from the *in situ* and dilution protocols. **a:** HiCRep shows that *in situ* Hi-C matrices are more reproducible than dilution matrices (*p* < 0.0005). **b,c:** TAD set reproducibility according to JI (*p* = 0.02703) and TADsim (*p* = 0.1547) shows that protocol choice has less of an impact on reproducibility of TAD sets than full Hi-C matrices. **d,e,f:** Comparisons of same cell type pairs generated by the same and different protocols. The background distribution is all comparisons of different cell types. Under all measures, there is a clear and statistically significant (*p* < 0.05) drop in similarity values of samples generated by different protocols compared to samples generated by the same protocol.

### Comparing *in situ* and dilution samples

In order to study whether both *in situ* and dilution protocols result in the same structures, we compared samples across protocols. Among pairs of the same cell type, mixed protocol pairs, where one sample came from *in situ* and one from dilution, were consistently statistically significantly less similar than the single protocol pairs, in which both samples came from the same protocol (Figures 3d,e,f, HiCRep: *p* = 0.0003423, JI: *p* = 0.03967, TADsim: *p* = 0.02002). The chromosomal structures identified from these two protocol variants are therefore not entirely consistent, although pairs from the same cell type still showed more similarity than pairs of different cell types, even among mixed protocol pairs (HiCRep: *p* = 3.436 × 10^−15^, JI: *p* = 1.2645 × 10^−9^, TADsim: *p* = 0.006010). We observe a similar trend across all non-replicate pairs as well (Figure S6). Overall, we observe some structural differences induced by the protocol variations, but not enough to obscure the general similarities expected from samples of the same cell type.

### Restriction enzyme choice has minimal impact on TAD sets

By comparing samples from the same lab of the same cell type, generated with a different restriction enzyme, we see no significant variation in similarity measures induced by restriction enzymes, as shown in Figure 4. As expected, the pairs of the same cell type with a different restriction enzyme tend to be more structurally similar than the background distribution, which includes all 2333 other pairwise comparisons. The choice of restriction enzyme does not appear to be a significant source of technical variation in measurements of chromosome structure, as both Hi-C matrices and TAD sets appear robust to this experimental variable.

**Figure 4:**
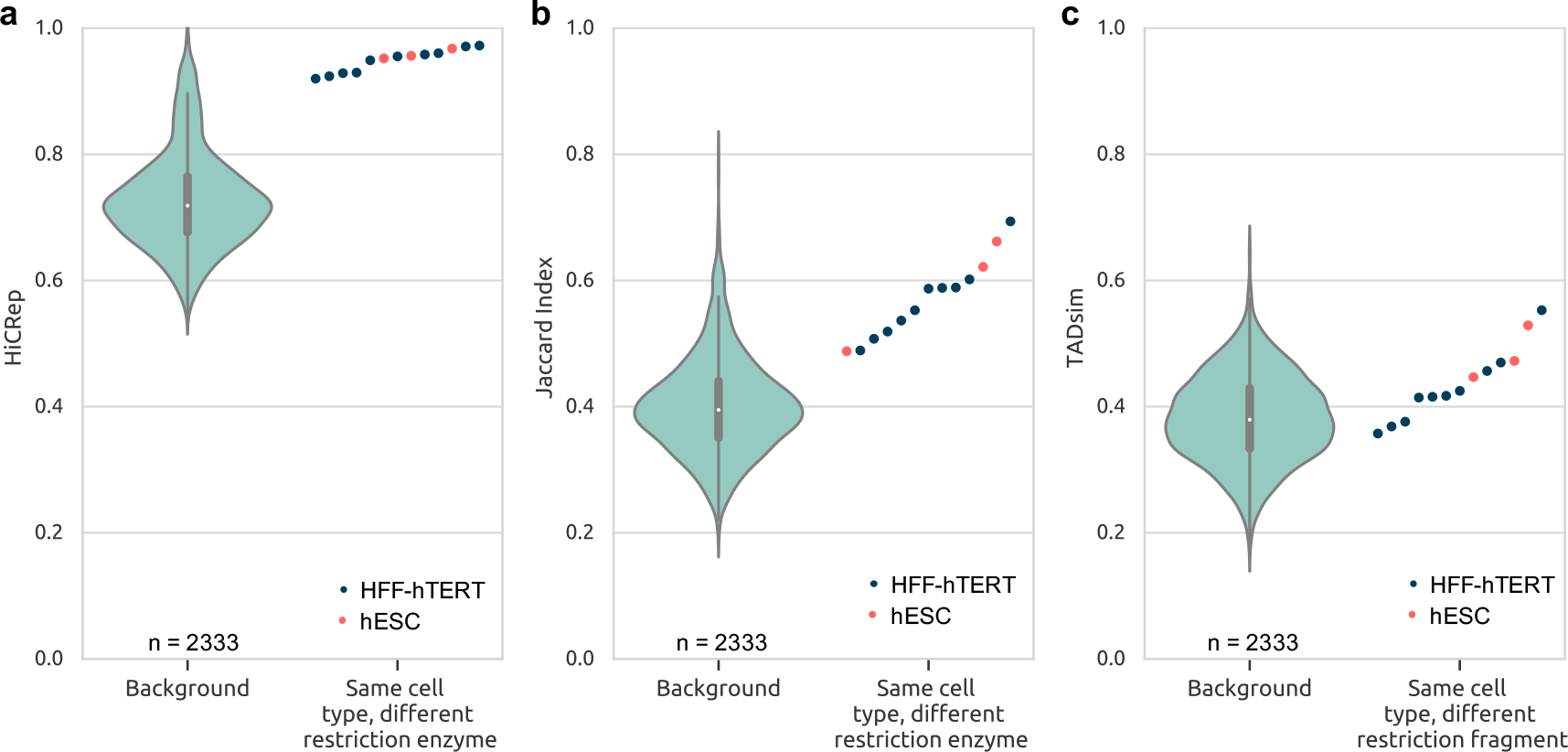
Measurements of structural similarity across the use of different restriction enzymes. The scattered points represent the similarity of a pair of samples of the same cell type (grey is hESC, red is HFF-hTERT), generated by using different restriction enzymes. The violin plot shows the distribution of all other non-replicate cmoparisons. As expected, the points that differ only in restriction enzyme are largely more similar than the background, suggesting that this choice is not a significant source of technical variability.

### TAD variation induced by lab-specific differences

Across all of our data, we see no pattern of elevated structural similarity among samples from the same lab (Figure 5a, JI and TADsim heat maps can be seen in Figures S7 and S8). A comparison of pairs of the same cell type from different labs shows that these pairs are generally more similar than non-replicate pairs, with similarity values near those of replicate pairs (Figures 5b,c,d). Consistent with the protocol-driven variation described above, the three lowest pairwise scores for IMR90 in both JI and TADsim are the three mixed protocol comparisons; all other points represent pairs generated by the same protocol. Chromosome structure seems to be robust to the variability across experimental labs.

**Figure 5:**
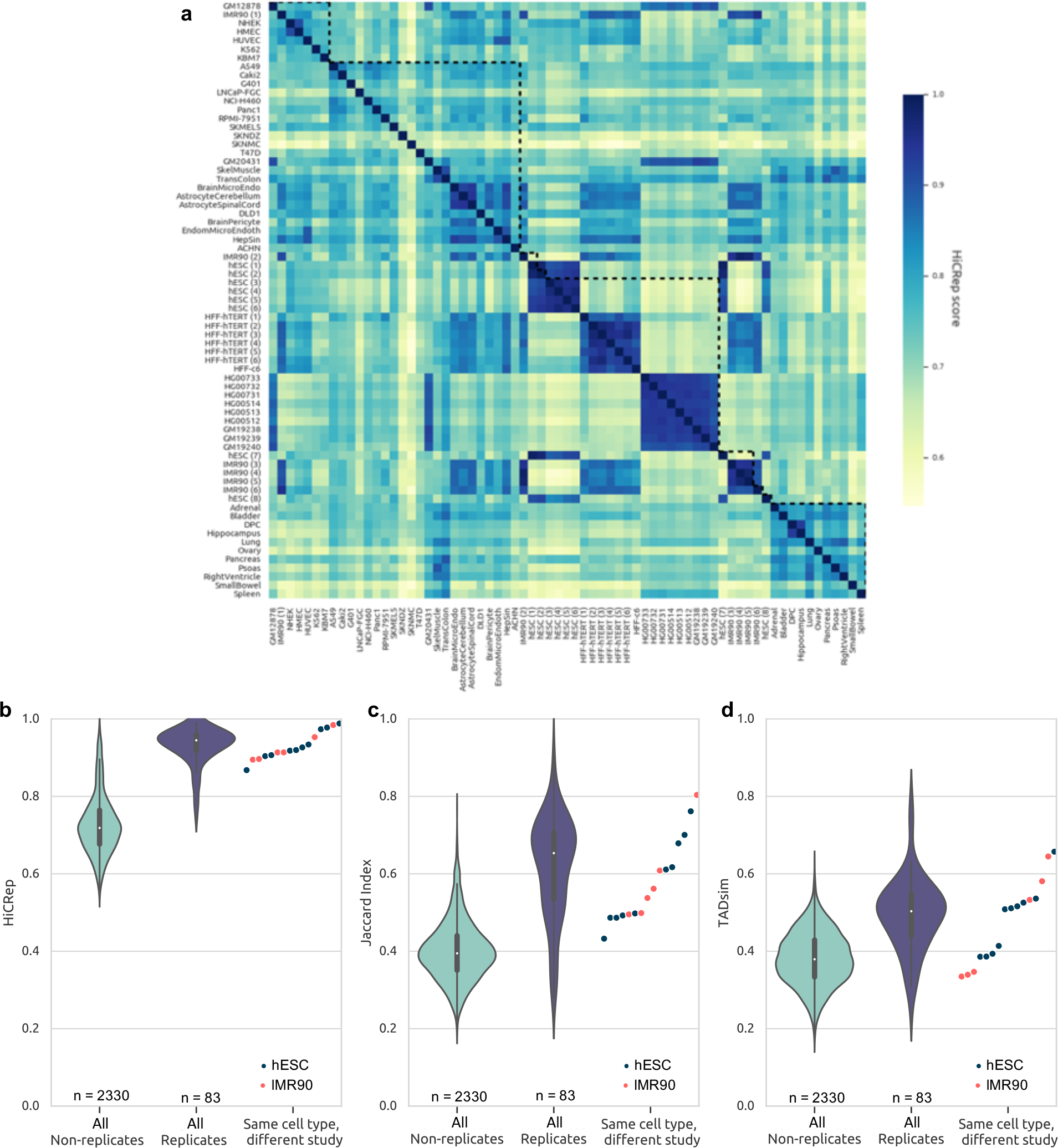
Quantifying variation across samples from different labs. **a** Summary of all 2346 pairwise sample comparisons as a heatmap of HiCRep scores. The dotted lines outline groups of samples from the same study. **b,c,d**: The effects of lab-specific variation on chromosome structure measurements. The points represent similarity scores of the same cell type (red for IMR90, grey for hESC) from different studies. These can be compared to the distribution of non-replicate pairs and that of replicate pairs, showing that samples from different labs achieve similarity values near those of replicate pairs.

## Discussion

We have demonstrated that cell or tissue type, rather than individual or genetic difference, appears to be the greatest driver of biological variation in TAD structures and Hi-C matrices, confirming and quantifying the likely biological importance of TADs. However, between replicates, TAD structures are shown to share only 60% of their boundaries, suggesting that chromosome structure is not a static feature, but remains variable even in identical cell populations. Contrary to previous claims that TADs are highly conserved, we note significant TAD variability across human samples. We observe elevated similarities between samples of the same cell type, suggesting that TAD structures are likely correlated with cellular function, rather than individual genetics. The largest differences due to technical variations appeared in comparing structures generated through *in situ* or dilution protocols, while lab-specific differences and restriction enzyme choices had a smaller impact on the resulting similarities of Hi-C measurements.

In order to maximize the number of samples analyzed in this work, all comparisons were performed at a fairly low resolution of 100kb, so structural features that would be clearer at higher resolution may have been overlooked. A few observations noted in this work are consistent with previous smaller-scale studies of higher-resolution matrices. In particular, the similarities between replicates that we observed were consistent with those found in Dixon *et al.* [11] and Rao *et al.* [35], though much higher than those reported in Forcato *et al.* [16] due to the different pre-processing methods used in each study. Using our pre-processing pipeline on the same data at the same resolution (40kb) as Forcato *et al.* [16] returned results consistent with our 100kb results. The higher-than-random similarity between pairs of different tissue types was also found by Schmitt *et al.* [39], but our quantification of this similarity suggests significant variability rather than extensive conservation between tissue types.

There are still relatively few available Hi-C data sets compared with other genomic analyses, and many of the observations made here would be strengthened with more samples or with confirmation through single-cell Hi-C. In particular, more trios from other cell types would help to confirm whether there is truly no elevated similarity in genetically related individuals, or whether this conclusion was specific to the blood lymphocyte samples studied in this work. As more single-cell Hi-C data becomes available and analysis methods improve, cell-to-cell variations in chromosome structure will be easier to assess, and we will be able to determine whether these population trends hold within individual cells.

All samples studied here were processed from sequencing reads to Hi-C matrices through the same pre-processing pipeline, and all TADs were computed using Armatus [15]. These choices may have influenced the trends we observed in this work, because different pre-processors, Hi-C normalization methods, or TAD callers could result in different patterns in the resulting structural measurements. The consistencies with previous work using different methods for each of these steps suggests that they did not have a major effect, but more study is needed to assess the overall robustness of Hi-C measurements to these processing choices. Additionally, there may be other possible comparison methods for Hi-C matrices and TAD sets, which may or may not agree with the three measures used here.

Further study of the structural differences across cell types may lead to insights into the mechanisms of chromosome structure. These comparison techniques could also be used to determine the differences between chromosome structures in healthy and diseased cells and could point to the locations of structural changes that are present across diseased cells. There is already significant evidence of structural abnormalities in many diseases (review, [25]). Additional systematic, genome-wide analyses of TAD structures could increase our understanding of a range of human diseases. Here, we have taken the first step towards systematically quantifying, at a large scale, the extent of TAD structure variability.

## Conclusions

This work compares Hi-C data and TAD structures from nine studies using three different measures, in order to identify trends in the variables controlling chromosomal structural similarity. We observe that even replicates display a certain amount of variability in chromosome structure. Chromosome structure appears most conserved within cell types and tissue types and not influenced more strongly by genetic similarity or differences across individuals. Differences in the cross-linkage step of the Hi-C protocol can induce variation in the resulting Hi-C and TAD measurements, but they seem robust to both lab-specific differences and choice of restriction enzyme.

## Methods

### Data

A total of 76 human Hi-C samples were processed from sequencing reads (.fastq files) downloaded from various publicly available sources (Sequence Read Archive (SRA) [23], ENCODE [42], Gene Expression Omnibus (GEO) [3], or 4DN portal [10]). Normalized Hi-C matrices were computed from the reads through the HiC-Pro pipeline [40], and each sample was tested for quality at 100kb resolution. Using the criteria suggested by Ay and Noble [1] and Rao *et al.* [35] (at least 80% of all bins must contain more than 1000 contacts), we found 7 experiments which could not be analyzed at 100kb resolution or less (Table S1), leaving 69 human Hi-C data sets (137 including all replicate samples) representing 52 unique cell types or biological sources from 9 studies. The details of these experiments, including accession numbers, are found in Table 1. All samples were normalized using iterative correction and eigenvector decomposition (ICE) [20], and all analyses presented here were performed at 100kb resolution. For analyses that do not explicitly compare replicates, all aligned reads from each replicate of a sample were merged and processed into a single combined Hi-C matrix for optimal data quality.

From the Hi-C matrices, TADs were computed using Armatus [15], a commonly used method for identifying TADs efficiently. Armatus takes one parameter *γ*, which controls the expected TAD size. For every sample and chromosome, Armatus was run with *γ* values ranging from 0 to 1 at intervals of 0.1, and the *γ* value was chosen to ensure a distribution of TADs with median as close as possible to the expected median TAD size of 880kb [4].

### Comparison measures

In order to comprehensively compare chromosome structures, we use three different measures: HiCRep [45], Jaccard Index (JI), and TADsim [38]. HiCRep measures similarity between Hi-C matrices directly, and both JI and TADsim compare similarity of predicted TADs. All three measures were computed on all 2346 pairs of non-replicate samples, in addition to all 83 replicate pairs.

HiCRep was designed to assess the reproducibility of replicates or the similarity of two Hi-C matrices. This measure uses a stratum-adjusted correlation coefficient to reliably compute a statistical similarity score between two Hi-C matrices, explicitly accounting for both the strong distance dependence found in Hi-C and the known domain structure [45]. This method returns a value that represents the overall similarity of the full Hi-C matrix, and distinguishes between replicate and non-replicate samples significantly better than simple correlation coefficients. We ran HiCRep on all intra-chromosomal matrices of our samples and averaged over all chromosomes to get a single value per cell type pair. HiCRep requires a smoothing parameter *h*, which was selected for each comparison according to the heuristic optimization procedure provided by the software, which chooses the minimum *h* value at which the score begins to converge. We allow a range of 0 to 3, which is expanded from the 0 to 2 range shown in the documentation example, to allow more options while maintaining computational efficiency.

The Jaccard Index (JI), a simple set similarity metric, is defined as the size of the intersection of two sets *A* and *B* divided by the size of the union of the sets: *JI*(*A, B*) = *|A ∩ B|/|A ∪ B|*. When comparing TADs, the two sets *A* and *B* represent the two lists of TAD boundaries, as used in Forcato *et al.* [16]. The resulting JI value is an easily interpretable number representing the fraction of shared boundaries between the two TAD sets.

While JI is an effective way to compare boundary locations, it does not take into account the total overlap between TAD interiors. We therefore also adopted a measure from Sauerwald and Kingsford [38], which presented a method to identify structurally similar regions between two TAD sets. We updated the original method to address some methodological artifacts and improve efficiency, as detailed in the Supplementary materials. The measure used here, which we will call “TADsim,” is the fraction of the genome covered by structurally similar regions identified by the method described in Sauerwald and Kingsford [38].

### Statistical comparisons

Distributions of similarity values under all three measures were checked for statistical significance through the Mann-Whitney *U* test, also called the Mann-Whitney-Wilcoxon (MWW) test. This nonparametric statistical test assesses the null hypothesis that a randomly selected value from one sample is equally likely to be less than or greater than a randomly selected value from the other sample. The alternative hypothesis can then be formulated as a randomly selected value from one distribution being likely to be greater than (or less than) a randomly selected value from the other distribution.

Without knowing the underlying distribution of structural similarity values, a nonparametric statistical test is required for all of our comparisons. The Kolmogorov-Smirnov two-sample test (KS test) is another commonly used nonparametric test, but it does not include any assessment of which distribution is greater than the other. The KS test is additionally sensitive not only to differences in the median or mean between two distributions, but any differences in their shapes, dispersion, or skewness as well. We therefore chose the MWW test for these analyses, given that we specifically are testing for the difference in relative magnitudes of the values in the distributions, rather than differences their overall shapes.

## Availability of data and material

All data used in this study are publicly available and listed, along with accession codes, in Table 1. The scripts to reproduce the analysis are available at https://github.com/Kingsford-Group/localtadsim/tree/master/analysis.

## Authors’ contributions

N.S. collected, processed, and analyzed all of the data. C.K. and N.S. designed the experiments and wrote the manuscript. A.S. conceived of and implemented all improvements to the original TADsim method. All authors contributed to the final manuscript.

## Acknowledgements

The authors would like to thank Dr. Mattia Forcato, Dr. Koustav Pal, and Dr. Chiara Nicoletti for valuable discussions.

