## Supplementary Material for "Analysis of the structural variability of topologically associated domains as revealed by Hi-C"

#### Modifications to TADsim method

The TADsim measure introduced in Sauerwald and Kingsford [38], uses the information theoretic variation of information (VI) metric [27] to measure the similarity between two sets of topologically-associating domains (TADs) and identify structurally similar sub-regions. We modify this method slightly to address a few concerns mentioned in the original work. All of the modifications are described below, and the code has been modified to reflect them (available at https://github.com/Kingsford-Group/localtadsim).

##### Correction: Local minima at TAD boundaries

The TADsim method calculates VI values for only those sub-intervals that begin and end at TAD boundaries. Although the VI formulation provides no theoretical guarantees of this, it seems intuitive that the minimum VI distance would occur largely at cluster boundaries. In order to test the validity of this intuitive understanding, we ran empirical tests counting the number of local minima that do not occur at TAD boundaries. It is mentioned in Sauerwald and Kingsford [38] that 97% of local minima occur at boundary points. However, we found that this result was due to a small bug in the code, and after correction we find that 100% of local minima occur at boundary points.

##### Hanging TADs

TADsim identifies structurally similar genomic regions by breaking the genome up into every possible interval that starts and ends at a TAD boundary in either TAD set, and testing each interval for statistically significant similarity of the TAD sets within them. One of the artifacts of this method is that the boundaries of these intervals may fall within a TAD, creating a false TAD boundary at the start of end of the interval, and the algorithm is unable to distinguish between true TAD boundaries and those imposed by the genome segmentation. We therefore define a "hanging TAD";

a TAD at the edge of an interval that has been truncated at less than 50% of its original length. Hanging TADs can appear to be a perfect match to a true TAD and will therefore be included in the interval identified by the method (an example of this can be seen in Figure S1, where the hanging TADs have been circled).

In order to avoid this, a preprocessing step has been added to the algorithm. Sub-intervals that include hanging TADs (as defined above) are removed from consideration before testing for statistical significance, guaranteeing that they will not be included in the output. Figure S2 shows the output for the same input set as Figure S1, after removing hanging TADs. Though there are fewer total intervals identified after this modification, the area covered by both outputs is essentially the same.

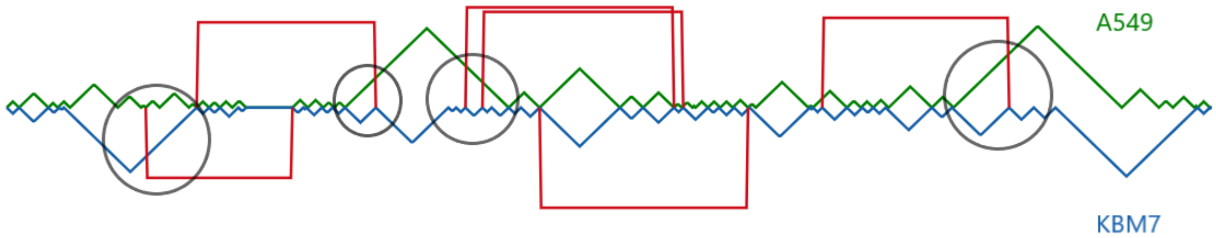


Figure S1: TADsim example. TAD sets from chromosome 18 are represented by triangles spanning each TAD, with TADs from A549 in green and from KBM7 in blue. The red brackets outline the significant, dominating intervals (regions of the genome covered by similar TAD structures) identified by the original TADsim method. Hanging TADs are shown by gray circles.

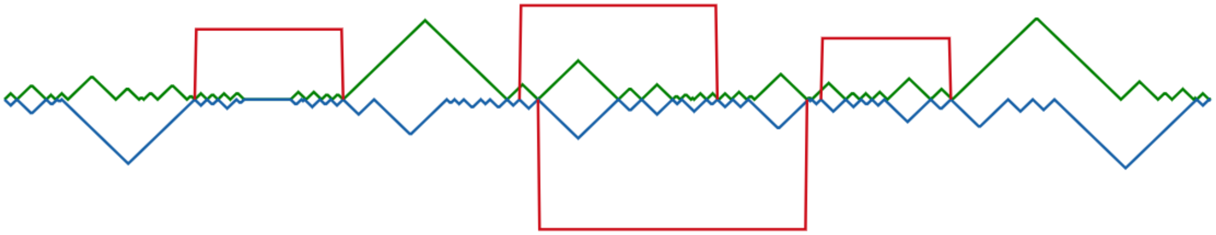

Figure S2: After removing hanging TADs. The same two TAD sets are shown, with the red brackets showing the output of the updated TADsim method which does not include hanging TADs in its output.

#### Adaptive shuffling

After computing the VI between TAD sets on every sub-interval, statistically significant sub-intervals are chosen by an adapted permutation test. For each sub-interval, one TAD set is fixed and TADs from the other set are randomly permuted 1000 times, calculating the VI at each permutation. Depending on the number of TADs being shuffled, 1000 permutations is either excessive or far too small. In some cases, this meant that results would differ on different runs with the exact same input due to under-sampling artifacts.

In order to improve the robustness of our results, we have modified the method to continue randomly permuting TADs until the p-values converge. We define convergence as the point where the p-values for 5 consecutive iterations are equal up to at least 5 decimal places. Though this approach appears more computationally intensive, in practice, it does not take more than twice as much time as the fixed 1000 shuffles. This is because, for very small numbers of TADs, convergence is reached very quickly and requires significantly fewer than 1000 permutations, somewhat balancing out the instances in which more than 1000 permutations are required.

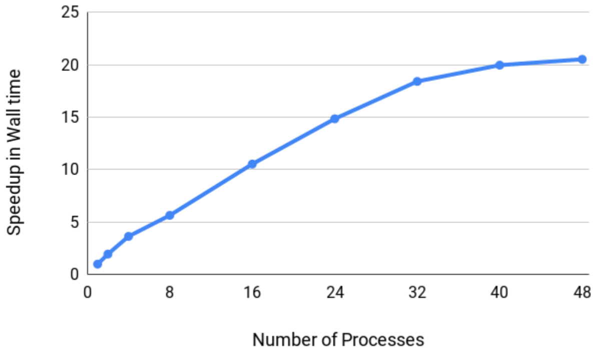


Figure S3: Speedup resulting from various code optimizations for tests ran on chromosome 1 of A549 and Caki2. The speedup is initially directly proportional to the number of processing cores used, but the speedup plateaus once the number of processes exceeds the number of cores, potentially due to processor saturation.

#### Parallelism, concurrency, and memory optimization

One of the challenges with the original implementation of TADsim was efficiency. Among all of the subroutines, the permutation testing is the most time intensive, taking up to several hours depending on the number of TADs in the input sets. Thus, we modified this method to make it

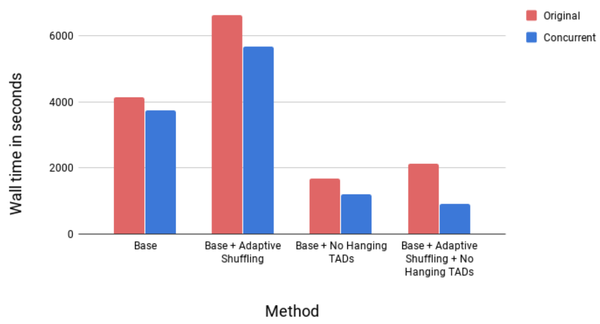


(a) Method changes

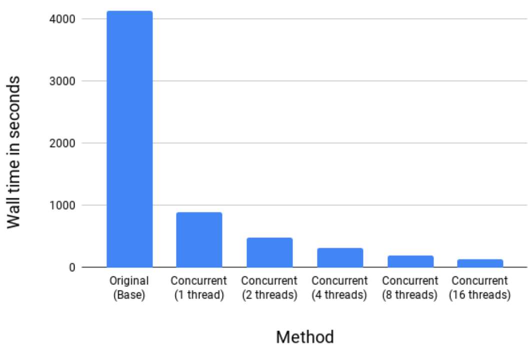


(b) Parallelization

Figure S4: Plots of the effects of methodological changes and parallelization on runtime.

both concurrent and memory optimized. In the current implementation, each execution can spawn one or more processes with each process having 1 or more threads. The independence of each permutation in the p-value calculation sub-routine allows for high concurrency that is maximized when each process has only one thread. The speedup is then directly proportional to the number of processing cores. However, once the process count becomes more than the number of cores there is only a minor improvement in the speedup which may be attributed to processor saturation (see Figure S3). The tests were done on a 24-core/48-thread 2.6 GHz Intel Xeon E5-2690 machine.

##### Execution Time

Each of the method changes described above affect the overall execution time of TADsim. For example, while adaptive shuffling often leads to an increase in the runtime, removing hanging TADs greatly reduces it, as shown in Figure S4a. To identify the effects of each of the functions we ran 22 random tests, 1 for each chromosome with random cell types on a 24-core/48-thread 2.6 GHz Intel Xeon E5-2690 machine. Figure S4b shows the runtime comparison of the original program against the modified program with different thread counts.

### Supplementary data & figures

| Cell type | Description | Replicates | Res frag | Protocol | Accession(s) | Citation |
|---|---|---|---|---|---|---|
| SJCRH30 | rhabdomyosarcoma fibroblast | 2 | HindIII | dilution | ENCLB379VAF, ENCLB821TDJ | [42] |
| GM06990 | blood lymphocyte | 1 | HindIII | dilution | SRR027956, SRR027957, SRR027958, SRR027959 | [24] |
| K562 | chronic myeloid leukemia | 1 | HindIII | dilution | SRR027962, SRR027963 | [24] |
| HeLa-S3 | cervix adenocarcinoma epithelial | 2 | HindIII | dilution | ENCLB693EVR, ENCLB696DUT | [42] |
| HepG2 | hepatocellular carcinoma epithelial | 2 | HindIII | dilution | ENCLB022KPF, ENCLB625TGE | [42] |
| hESC | human embryonic stem cell | 2 | HindIII | *in situ* | GSE70181 | [30] |
| hESC | human embryonic stem cell | 2 | HindIII | dilution | GSE70181 | [30] |

Table S1: Hi-C samples downloaded and processed that could not be analyzed at 100kb resolution

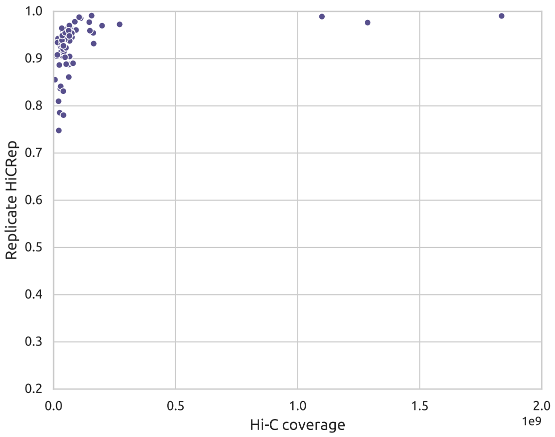


(a) HiCRep; Spearman $\rho = 0.6378$

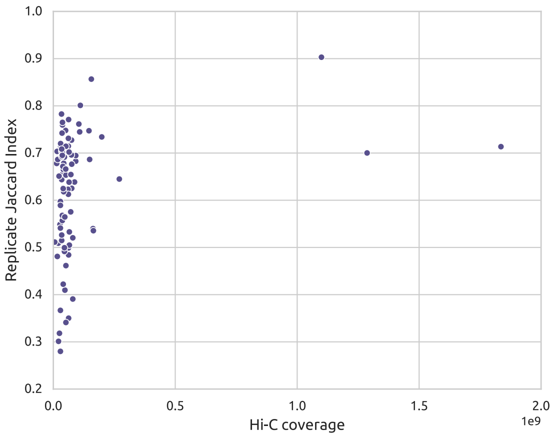


(b) JI; Spearman $\rho = 0.2407$

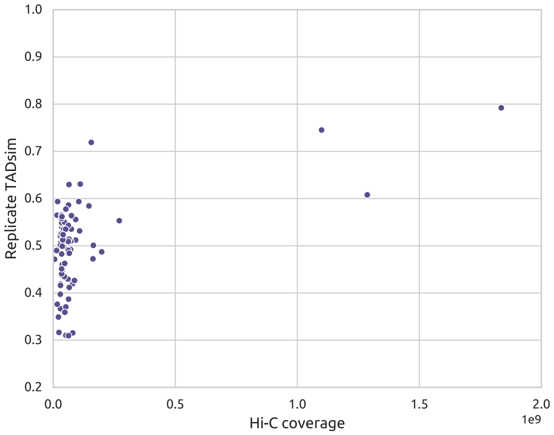


(c) TADsim; Spearman $\rho = 0.2310$

Figure S5: **Reproducibility versus Hi-C coverage.** Reproducibility is quantified by the similarity value of each replicate pair. We compute a contact count for each replicate sample by adding all non-normalized contacts on all intra-chromosomal matrices, and Hi-C coverage is defined as the smaller of the two contact counts for the replicate samples being compared. Across all three measures, especially the two quantifying TAD reproducibility, there is a low correlation with coverage though very high coverage experiments tend to have high reproducibility.

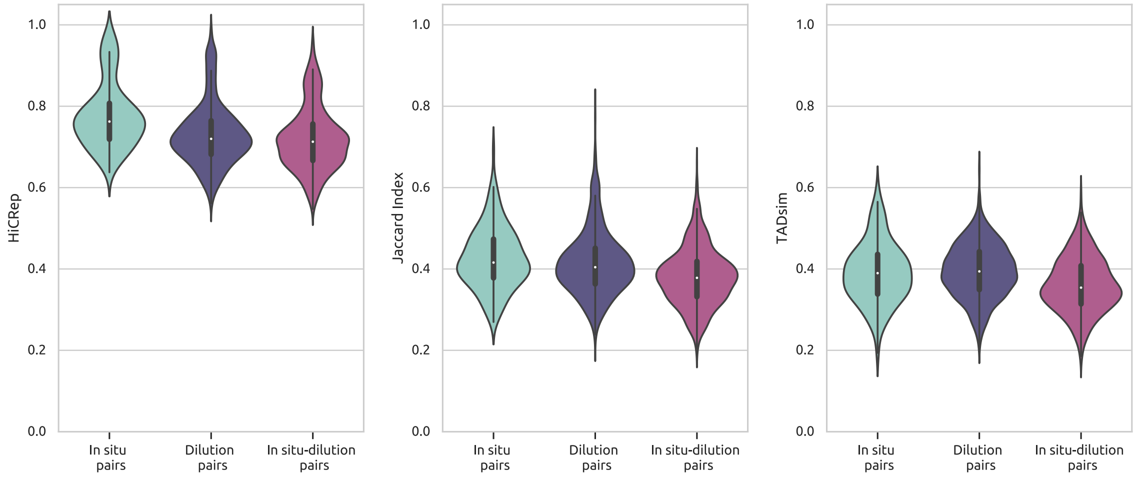


(a) HiCRep (b) Jaccard Index (c) TADsim

Figure S6: All pairwise comparisons, divided into three distributions: pairs in which both samples came from an *in situ* experiment, pairs in which both came from a dilution experiment, and pairs in which one sample was dilution and one was *in situ*. Under all measures, the pairs containing samples from the two different protocols were statistically significantly lower than pairs from the same protocol ($p < 0.01$), demonstrating that samples produced through the different protocols have lower similarity due simply to this technical variation rather than meaningful biological differences.

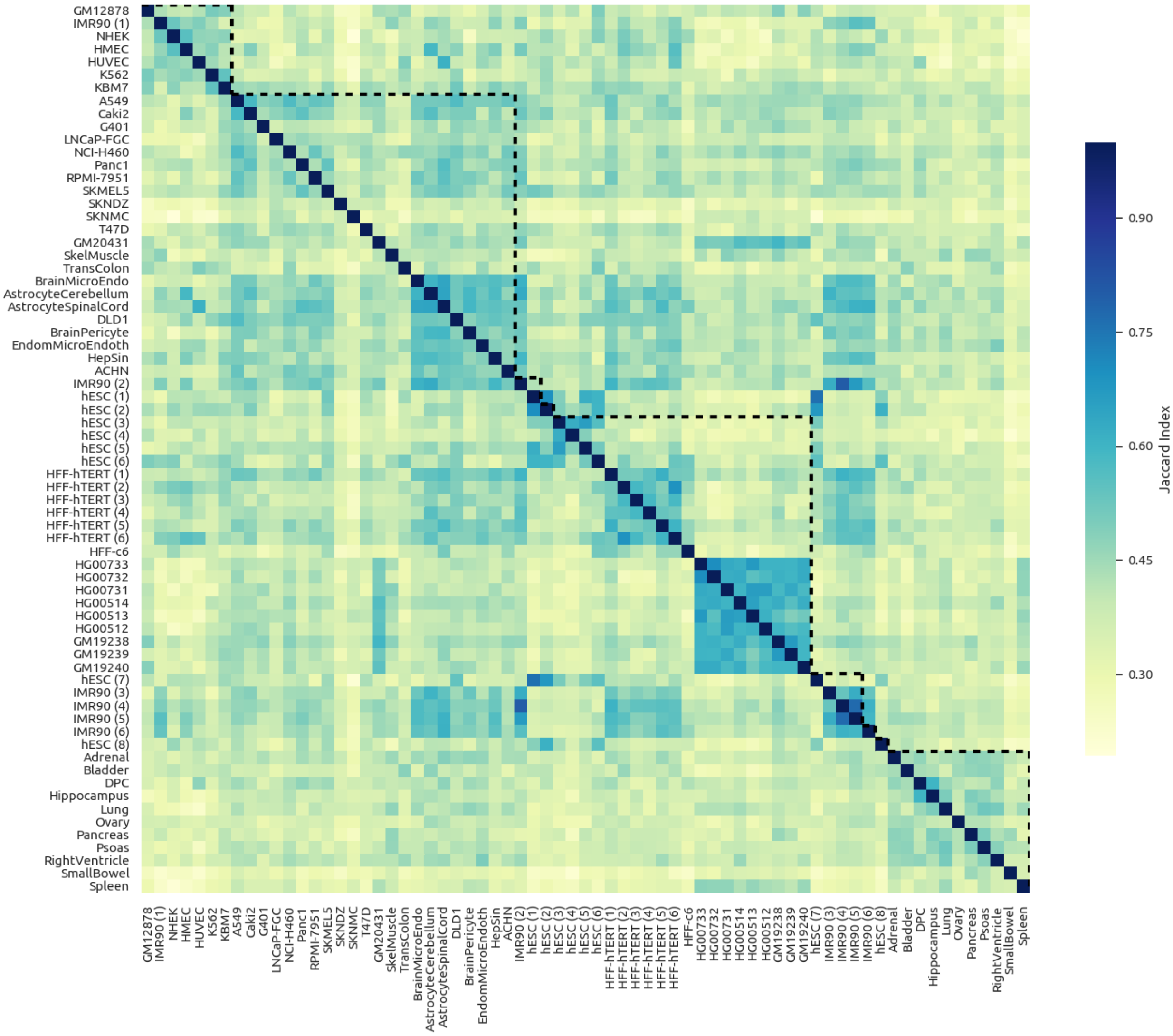


Figure S7: Summary of all 2346 pairwise sample comparisons as a heatmap of JI values. Dotted lines mark the samples that came from the same study. We see no systematic elevation in similarity values of intra-lab comparisons. This suggests that lab-specific variations do not significantly impact the similarities of TAD sets.

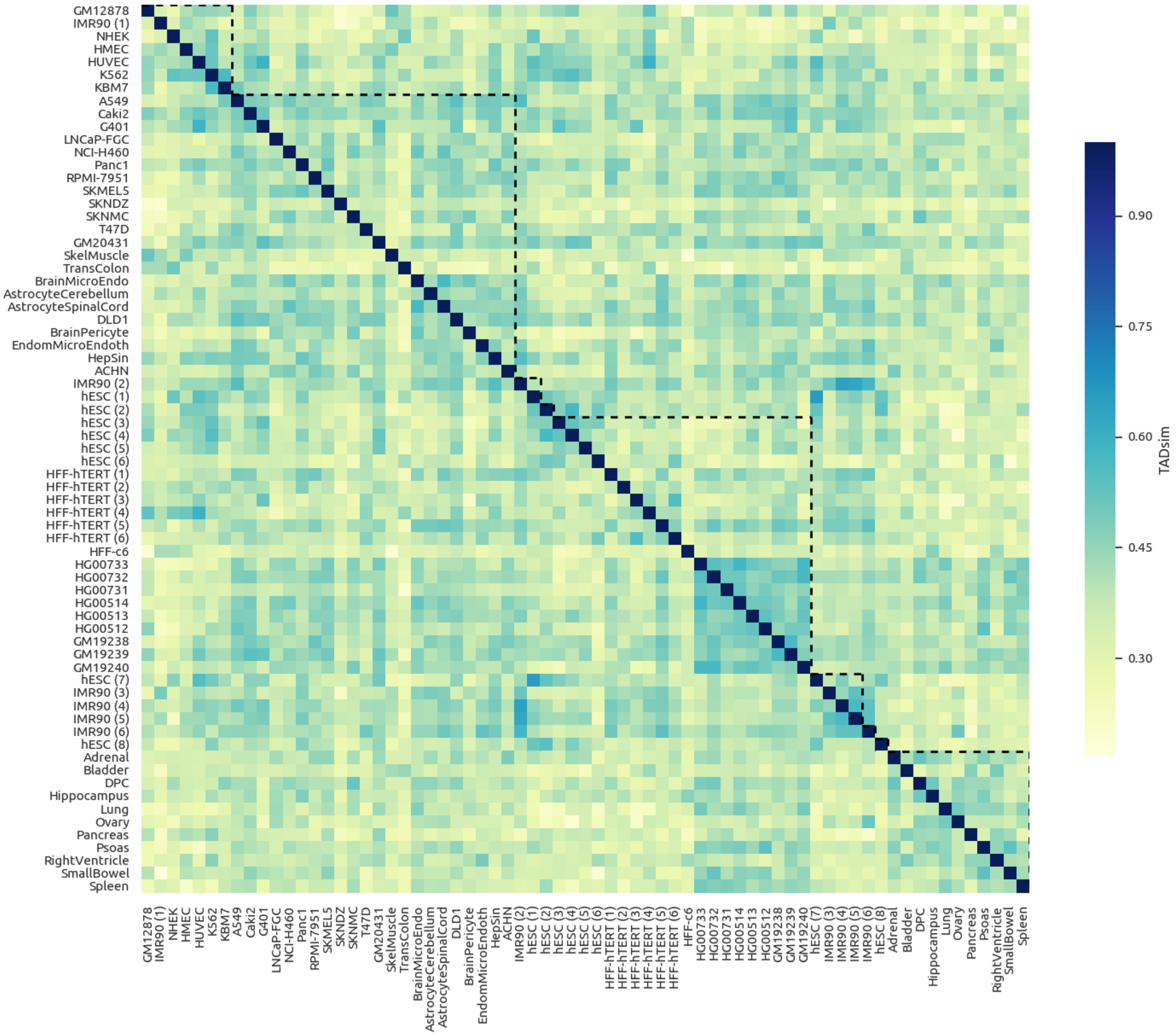


Figure S8: Summary of all 2346 pairwise sample comparisons as a heatmap of TADsim values. Dotted lines mark the samples that came from the same study. We see no systematic elevation in similarity values of intra-lab comparisons. This suggests that lab-specific variations do not significantly impact the similarities of TAD sets.
